# Competitive modulation of Kv1.2 gating by LMAN2 and Slc7a5

**DOI:** 10.1101/2024.07.25.605230

**Authors:** Damayantee Das, Shawn M. Lamothe, Anson Wong, Victoria A. Baronas, Harley T. Kurata

**Author notes:** **Corresponding author:** Harley T. Kurata, Current address: Dept. of Pharmacology, Alberta Diabetes Institute, University of Alberta, 9-70 Medical Sciences Building, Edmonton, AB, T6G 2H7, Canada.

## Abstract

Kv1.2 is a prominent ion channel in the CNS, where it regulates neuronal excitability. Kv1.2 structure and function are well understood, but there is less consensus on mechanisms of regulation of Kv1.2 and other potassium channels by auxiliary proteins. We identified novel regulators of Kv1.2 by a mass spectrometry approach. The neutral amino acid transporter Slc7a5 causes a dramatic hyperpolarizing shift of channel activation. In contrast, LMAN2 is a recently identified candidate regulator that has the opposite effect on gating: large depolarizing voltages are required to activate Kv1.2 channels co-expressed with LMAN2. In this study, we characterized the functional interaction between LMAN2 and Slc7a5 on Kv1.2 gating properties and identified key structural elements that underlie sensitivity to each regulator. When LMAN2 and Slc7a5 are expressed together, Kv1.2 activation exhibits a bi-modal voltage-dependence, suggesting two distinct populations of channels regulated either by LMAN2 or Slc7a5, but not both. Using a Kv1.2:1.5 chimeric approach, we identified specific regions between the S1 to S3 segments of the voltage sensing domain (VSD) that are distinct for either Slc7a5 or LMAN2 sensitivity. By replacing either segment with sequence from Kv1.5, modulation by the corresponding regulator was selectively abolished. These results suggest that Slc7a5 and LMAN2 compete for interaction with the Kv1.2 voltage sensor, leading to complex voltage-dependence of channel activity when both regulators are present in the cell.

## INTRODUCTION

Voltage-gated potassium (Kv) channels are the most diverse ion channel family, and influence electrical activity in most excitable cells. Kv1.2 is thought of as a prototypical delayed rectifier channel, and is prominently expressed in the central nervous system where it regulates action potential properties including firing frequency, repolarization, and threshold (Kole *et al*., 2007; Trimmer, 2015). In the context of understanding basic ion channel gating mechanisms, Kv1.2 has been a widely used model for interpreting structure-function studies of Kv channels, as it was the first mammalian Kv channel used to generate atomic resolution structures (Long *et al*., 2007). In addition to this historical importance as a template for relating structure and function, it is often overlooked that Kv1.2 is prone to powerful modulation. For Kv1.2 in particular, widely variable gating properties have been reported, depending on the expression system and recording conditions. Therefore, these channels are susceptible to prominent regulation to cellular factors that are extrinsic to the channel (Grissmer *et al*., 1994; Kerr *et al*., 2001; Rezazadeh *et al*., 2007; Baronas, McGuinness, *et al*., 2015; Baronas, Yang, *et al*., 2015; Baronas *et al*., 2017) .

Among closely related (*Shaker*/Kv1 subfamily) channels, Kv1.2 also stands out in terms of the severity of phenotypes that arise with human mutations or in knockout organisms (Brew *et al*., 2007; Syrbe *et al*., 2015; Masnada *et al*., 2017; Nilsson *et al*., 2022). Severe outcomes associated with Kv1.2 mutations include epileptic encephalopathy, intellectual disability, developmental disorder and episodic ataxia (Chen *et al*., 2016; Corbett *et al*., 2016; Helbig *et al*., 2016). Moreover, Kv1.2 knockout mice exhibit 100% lethality within a few weeks of birth due to generalized seizures (Brew *et al*., 2007). These severe neurological outcomes are consistent with important and potentially specialized roles in the CNS, where Kv1.2 is found in most neurons and may influence diverse aspects of neuronal function (Wang *et al*., 1994; Rhodes *et al*., 1996; Sheng and Kim, 1996; Pinatel and Faivre-Sarrailh, 2020).

While biophysical studies of Kv channels most often investigate fundamental properties of homomeric channels, in physiological settings most Kv channels likely assemble as heteromers with other subfamily members. In the CNS, Kv1.2 frequently assembles into heteromeric channels with Kv1.1 and/or Kv1.4 (Isacoff *et al*., 1990; Sheng *et al*., 1993; Wang *et al*., 1993, 2013; Dodson *et al*., 2002; Ovsepian *et al*., 2013).

Various proteins have been implicated in the regulation of Kv1 channels. The most well-characterized regulators of Kv1.2 and other Kv1 channels are the Kvβ subunits, which promote surface maturation and can also impart N-type inactivation (Rettig *et al*., 1994; Shi *et al*., 1996), and their association with Kv1.2 has been described with atomic resolution structures (Gulbis *et al*., 1999, 2000). However, in terms of other potential regulatory proteins, there is often not strong consensus or clarity in terms of mechanism or physiological roles. Function and surface expression of Kv1.2 has been reported to be influenced by various regulators including M1 muscarinic acetylcholine receptors (suppression of the channel in a tyrosine-phosphorylation dependent manner)(Huang *et al*., 1993), secretin, or signaling pathways that influence cAMP levels (Nesti *et al*., 2004; Connors *et al*., 2008; Williams *et al*., 2012).

Another potential example is the actin-binding protein cortactin, which is reported to bind to Kv1.2 in specialized axonal segments and is modulated by tyrosine phosphorylation (Hattan *et al*., 2002). Kv1.2 may also be modulated by phospholipids, as PIP2 depletion or phosphatidic acid enrichment have been reported to shift the voltage dependence of activation and current level (Rodriguez-Menchaca *et al*., 2012; Hite *et al*., 2014). Axonal ADAM23 promotes accumulation of Kv1 complexes at the juxtaparanodal region of neurons (Kozar-Gillan *et al*., 2023), and HDAC2 has been found to regulate Kv1.2 in DRG neurons to mediate neuropathic pain (Li *et al*., 2019). Within this complex landscape of potential regulatory proteins, there is not a strong understanding of the associated gating effects, structural determinants of sensitivity, interactions between regulators, or consensus on biological roles.

We have recently reported additional regulatory proteins that cause extremely prominent gating effects when co-expressed with Kv1.2, including the neutral amino acid transporter Slc7a5, and the transmembrane lectin LMAN2 (Baronas *et al*., 2017, 2018; Lamothe *et al*., 2020, 2024). These diverse regulators generate a remarkably wide range of voltage-dependence in Kv1.2, with V_1/2_ s ranging between roughly -60 mV and +60 mV. However, we lack a complete understanding of the underlying mechanisms involved in this uniquely large dynamic range of modulation. Since the gating effects of LMAN2 and Slc7a5 are so powerful, we have sought to use these proteins as models to begin to investigate structural determinants that underlie Kv1.2 sensitivity to modulation, and also potential ways that different modulators may functionally interact in terms of Kv1.2 activity. We show here that Slc7a5 and LMAN2 act via distinct mechanisms, and we identify structural elements in the voltage sensor domain that can specifically abolish sensitivity to Slc7a5 or LMAN2 modulation. Finally, we observe that varying the relative expression level of LMAN2 and Slc7a5 can lead to a wide dynamic range of different gating behaviors, suggesting that competition between these and potentially other regulators can strongly influence Kv1.2 gating.

## MATERIALS AND METHODS

### Cell culture

Mouse L(tk-) fibroblast cells (ATCC CCL-1.3), referred to throughout as LM cells, were used for whole-cell patch clamp studies. HEK293 cells (ATCC) were used for Western blot experiments. Cell culture was maintained at 37°C in a 5% CO_2_ incubator in DMEM supplemented with 10% FBS and 1% Penicillin/Streptomycin. Cells were transfected with cDNA using jetPRIME transfection reagent (Polyplus). For electrophysiology, the cells were plated into 12 wells and replated onto coverslips 6 h after transfection. Recordings were done 24 h post-transfection. For Western blot, cells were seeded into 12 well plates, transfected the following day, then lysed 72 hours post-transfection.

### Plasmid constructs

cDNAs were expressed in the pcDNA3.1(-) vector (Invitrogen). Fluorescent proteins mCherry and EGFP were fused with Slc7a5 and LMAN2 respectively using standard PCR and compatible restriction digestion and ligation. The Kv1.2-Kv1.5 chimeras (S1 and S2-S3L) were constructed by amplifying the S1 and S2-S3L segments of Kv1.5, and then replacing the corresponding segments in Kv1.2 by two-step overlapping PCR, as previously described (Lamothe *et al*., 2020). The Kv1.2-Kv1.5 (S1-S3) chimera was constructed by amplifying an N-terminal fragment from a previously described Kv1.2:Kv1.5 chimera (Lamothe *et al*., 2020), comprising N-terminal sequence of Kv1.2 up to residue 153 (intracellular boundary of S1), followed by sequence from Kv1.5. A second fragment from Kv1.2, comprising the S3 segment to the C-terminus. These fragments were combined by overlapping PCR and cloning into pcDNA3.1(-) using NheI and HindIII, leading to a chimera that replaces residues 153-259 of Kv1.2 with corresponding sequence from Kv1.5. Generation of this chimera was previously described (Lamothe *et al*., 2024). Constructs were verified using diagnostic restriction digestion, followed by Sanger sequencing. For electrophysiology experiments, channel and LMAN2 constructs were expressed in a 1:4 ratio, while the channel to Slc7a5 was expressed in 1:0.5 ratio. The total amount of DNA for the experiments was held constant using either GFP plasmid or mCherry or empty pcDNA3.1 plasmid for transfection.

### Electrophysiology

Patch pipettes were made from soda lime capillary glass (Fisher), using a Sutter P-97 puller (from Sutter instrument). While recording with standard internal solution, the tip resistance was between 1-3 MΩ. Recordings were filtered at 5 KHz, and sampled at 10 KHz, with manual capacitance compensation and series resistance compensation between 70 to 80%. The data was stored using Clampex software (Molecular Devices). The external (bath) solution composition was 135 mM NaCl, 5 mM KCl, 1 mM CaCl_2_, 1 mM MgCl_2_, 10 mM HEPES and the pH was adjusted to 7.3 using NaOH. The internal pipette solution composition was 135 mM KCl, 5 mM K-EGTA, 10 mM HEPES and the pH was adjusted to 7.2 using KOH. Chemicals were purchased from either Sigma-Aldrich or Fisher. DTT was dissolved in ddH_2_O then stored in -20℃ as a 1 mM stock solution then diluted to 200 µM working concentration on the day of experiment. Cells for patch clamp recordings were identified using co-transfected fluorescent proteins or fluorescent protein tags fused to LMAN2 or Slc7a5. Simultaneous visualization of EGFP and mCherry fluorescence (to assess expression of EGFP-LMAN2 and mCherry-Slc7a5) was done using a multi-bandpass filter (Chroma 86021 filter set with excitation wavelengths of 470/30 nm, 565/55 nm; emission wavelengths of 510/30 nm, 650/75 nm).

### Data analysis

Data are reported throughout as mean + SEM or box plots along with data points from individual cells (box plots depict the median, 25^th^ and 75^th^ percentile (box), and 10^th^ and 90^th^ percentile (whiskers)). Conductance-voltage relationships were fitted using a Boltzmann equation (Equation 1), where G is the normalized conductance, V is the voltage applied, V_1/2_ is the half-activation voltage, and k is a fitted value reflecting the steepness of the curve.

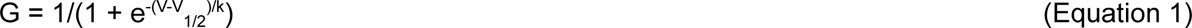

In some cases, conductance-voltage relationships were fit better with a two-component Boltzmann equation”

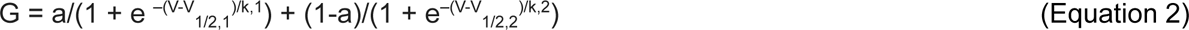

In equation 2, V_1/2,1_ and V_1/2,2_ are half-maximal voltages of each component, k_1_ and k_2_ are slope factors of respective components and a is the fraction.

Conductance-voltage relationships were fit for each individual cell, and the extracted fit parameters were used for subsequent statistical calculations.

### Western blot

Cell lysates from HEK293 were collected in NP40 lysis buffer (1% NP-40, 150 mM NaCl, 50 mM Tris-HCl) and 1% Protease Inhibitor cocktail (Sigma, P8340), 72 hours post-transfection. The samples were separated using 8% SDS PAGE gels and then transferred onto a nitrocellulose membrane using standard Western blot apparatus (Bio-Rad). Kv1.2 was detected using a mouse monoclonal Kv1.2 antibody (1:10,000 dilution; clone K14/16 75008; NeuroMab) and HRP-conjugated goat anti-mouse antibody (1:30,000 dilution; HS023; Applied Biological Materials). β-actin was used as a loading control and detected using the β-actin monoclonal antibody (1:10,000 dilution; GT5512; GeneTex). Chemiluminescence was detected using a SuperSignal West Femto Max Sensitivity Substrate (Thermo Fisher Scientific) and a ChemiDoc imaging system (BioRad).

## RESULTS

### Multimodal regulation of Kv1.2

Previous reports have described wide variability of Kv1.2 gating properties, depending on the expression system, recording conditions (including extracellular redox potential), and co-expression with regulatory proteins (Baronas *et al*., 2017, 2018; Abraham *et al*., 2019; Lamothe and Kurata, 2020; Lamothe *et al*., 2020). Under conventional recording conditions (with ‘ambient’ unbuffered redox potential), the reported V_1/2_ of Kv1.2 varies widely within and amongst studies, between -40 mV and +30 mV (Grissmer *et al*., 1994; Rezazadeh *et al*., 2007; Syrbe *et al*., 2015; Baronas *et al*., 2017; Abraham *et al*., 2019). We have reported that the voltage-dependence of Kv1.2 activation can be biased to positive voltages by application of reducing agents (Baronas *et al*., 2017) or co-expression with LMAN2 (Lamothe *et al*., 2024). In contrast, Kv1.2 gating is prominently shifted to extreme negative voltages when co-expressed with Slc7a5 (Figure 1B) (Baronas *et al*., 2018; Lamothe and Kurata, 2020; Lamothe *et al*., 2020). In confirmatory experiments in this study, we observed that co-expression of Slc7a5 with Kv1.2 causes a hyperpolarizing shift of the V_1/2_ of activation to -45.8 ± 3.5 mV, compared to 19.1 ± 6.8 mV for Kv1.2 alone (Figure 1B,C,D), consistent with previous reports (Baronas *et al*., 2018; Lamothe *et al*., 2020). In contrast, Kv1.2 exhibits a depolarizing shift of the voltage dependence of activation to a V_1/2_ of 48.3 ± 8.5 mV, when coexpressed with LMAN2, along with pronounced deceleration of the kinetics of activation (Figure 1B,E). Thus, Kv1.2 gating can be modulated over a broad voltage range, under the influence of co-expressed candidate regulatory proteins such as Slc7a5 or LMAN2.

**Figure 1.**
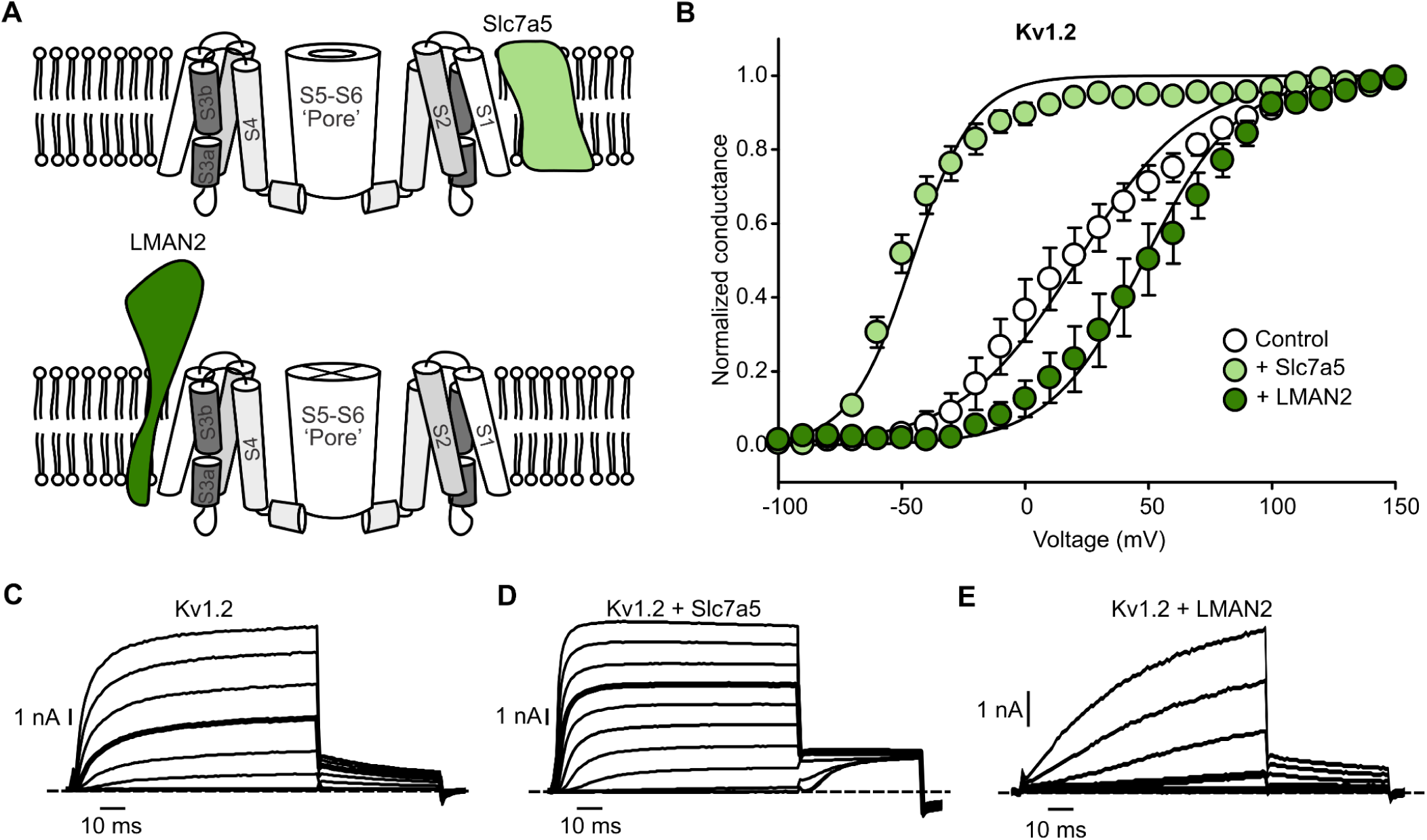
Multi-modal regulation of Kv1.2 generates wide variation of voltage-dependent gating. (A) Schematic model of Kv1.2 channel associated with candidate interactors Slc7a5 or LMAN2. (B) Conductance voltage-relationships for Kv1.2 co-expressed with Slc7a5 and with LMAN2, as indicated. Conductance-voltage relationships were generated by stepping between -100 mV and +150 mV (100 ms in 10 mV steps, -100 mV holding potential). Normalized tail current amplitudes were measured at -30 mV and fit with a Boltzmann function (mean ± SEM, Kv1.2 V_1/2_ = 19.1 ± 6.8 mV, k = 24.4 ± 1.2 mV, n=9; Kv1.2 + Slc7a5, V_1/2_ = -45.8 ± 3.5 mV, k = 14.3 ± 1.7 mV, n=10; Kv1.2 + LMAN2 V_1/2_ = 48.3 ± 8.5 mV, k = 20.0 ± 1.6 mV, n=7). (C,D,E) Exemplar patch clamp records from WT Kv1.2, Kv1.2 + Slc7a5 and Kv1.2 + LMAN2 as indicated. Voltage steps are shown in 20 mV intervals for clarity. Pulses to +40 mV have been highlighted in boldface for comparison between conditions.

### The voltage sensor controls Slc7a5 and LMAN2 sensitivity of Kv1.2

Unlike Kv1.2, Kv1.5 subunits are insensitive to both Slc7a5 and LMAN2 (Baronas *et al*., 2018; Lamothe *et al*., 2024). We used a chimeric strategy of substituting regions of Kv1.2 with homologous regions of Kv1.5 to investigate determinants of LMAN2 and Slc7a5 sensitivity. We first observed that a chimeric channel generated by replacing Kv1.2 sequence from the beginning of S1 until the intracellular side of S3, with corresponding sequence from Kv1.5 (Figure 2A), was insensitive to both Slc7a5 (Figure 2B,D) and LMAN2 (Figure 2B,E). Importantly, this Kv1.2-Kv1.5(S1-S3) chimera expressed alone exhibited very little cell-to-cell variability of voltage-dependent gating, with a V_1/2_ = 0.5 ± 3.3 mV, demonstrating that the voltage sensing domain is likely an essential determinant of regulation that leads to widely variable gating properties of Kv1.2 (Figure 2B,C). The voltage-dependence of the Kv1.2-Kv1.5(S1-S3) chimera was minimally affected by co-expression with either Slc7a5 (V_1/2_ of -7.5 ± 3.6 mV) or LMAN2 (V_1/2_ = -4.6 ± 3.7 mV) (Figure 2B, also summarized later for all chimeras tested).

**Figure 2.**
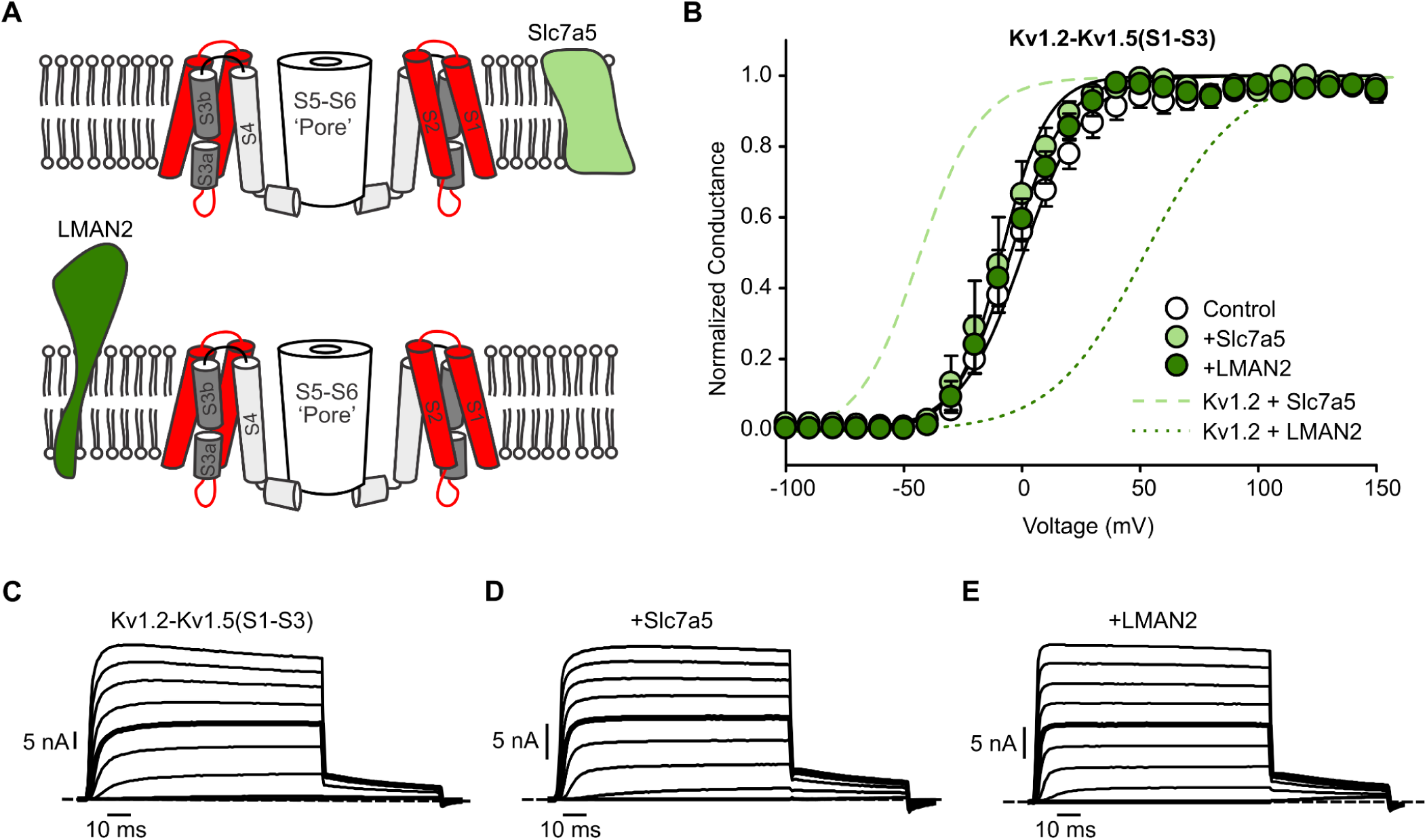
Extensive voltage sensor substitution abolishes dynamic voltage sensitivity in Kv1.2. (A) Schematic model of Kv1.2-Kv1.5(S1-S3) chimera channel along with candidate interactors Slc7a5 or LMAN2, with chimeric substitution of the Kv1.5 S1-S3 segment highlighted in red. (B) Conductance voltage-relationships for Kv1.2-Kv1.5(S1-S3) co-expressed with Slc7a5 or LMAN2 as indicated, using an identical protocol as Figure 1 (dashed lines are duplicated from Figure 1 for reference). Conductance-voltage fits were (mean ± SEM, Kv1.2-Kv1.5(S1-S3) V_1/2_ = 0.5 ± 3.3 mV, k = 15.0 ± 2.3 mV, n=7; Kv1.2-Kv1.5(S1-S3) + Slc7a5, V_1/2_ = -7.5 ± 3.6 mV, k = 11.1 ± 1.0 mV, n=6; Kv1.2-Kv1.5(S1-S3) + LMAN2 V_1/2_ = -4.6 ± 3.7 mV, k = 12.4 ± 1.5 mV, n=4). (C, D, E) Exemplar patch clamp records from Kv1.2-Kv1.5(S1-S3), Kv1.2-Kv1.5(S1-S3) + Slc7a5 and Kv1.2-Kv1.5(S1-S3) + LMAN2 as indicated. Voltage steps are shown in 20 mV intervals, and pulses to +40 mV are highlighted in boldface for comparison between conditions.

### Decomposition of LMAN2 and Slc7a5 sensitivity

We used smaller chimeric substitutions of Kv1.2 and Kv1.5 to highlight distinct determinants of sensitivity to Slc7a5 and LMAN2. We confirmed previous findings demonstrating that the S1 segment is an important determinant of Slc7a5 modulation of Kv1.2 and Kv1.1, as the Kv1.2-Kv1.5(S1) chimera resulted in abolished sensitivity to the Slc7a5-mediated hyperpolarizing shift (Figure 3B-D) (Lamothe *et al*., 2020). Instead, the Kv1.2-Kv1.5(S1) chimera was strongly biased towards a ‘slow’ gating phenotype, including slow activation kinetics (Figure 3C-E), and a relatively positive V_1/2_ of activation (Figure 3B). That is, these features generally resemble the slower mode of gating that is occasionally observed for WT Kv1.2 expressed alone (Baronas, McGuinness, *et al*., 2015; Baronas *et al*., 2017; Lamothe *et al*., 2024). It is noteworthy that considerable variability is retained in this chimera, in terms of the conductance-voltage behavior, leading to some divergence of the averaged data from a simple Boltzmann fit (Figure 3B). Despite this variability, in this case we have used single Boltzmann fits to simplify comparisons between conditions. Kv1.2-Kv1.5(S1) expressed alone exhibited a mean V_1/2_ of 34.9 ± 9.8 mV, whereas co-expression with Slc7a5 led to a V_1/2_ of 11.8 ± 6.9 mV, deviating significantly from the large hyperpolarized V_1/2_ that is observed from WT Kv1.2 co-expressed with Slc7a5 (shown as light green dashes in Figure 3B for comparison). This is the most important feature of this chimera - co-expression with Slc7a5 completely fails to replicate the large Slc7a5-mediated V_1/2_ shift observed in WT Kv1.2. Co-expression with LMAN2 did not have a prominent effect in this case (V_1/2_ of 26.2 ± 11.5 mV), although this is likely because the Kv1.2-Kv1.5(S1) chimera expressed alone is already strongly biased towards ‘slow’ gating properties that mimic LMAN2-dependent modulation (Figure 3B,E).

**Figure 3.**
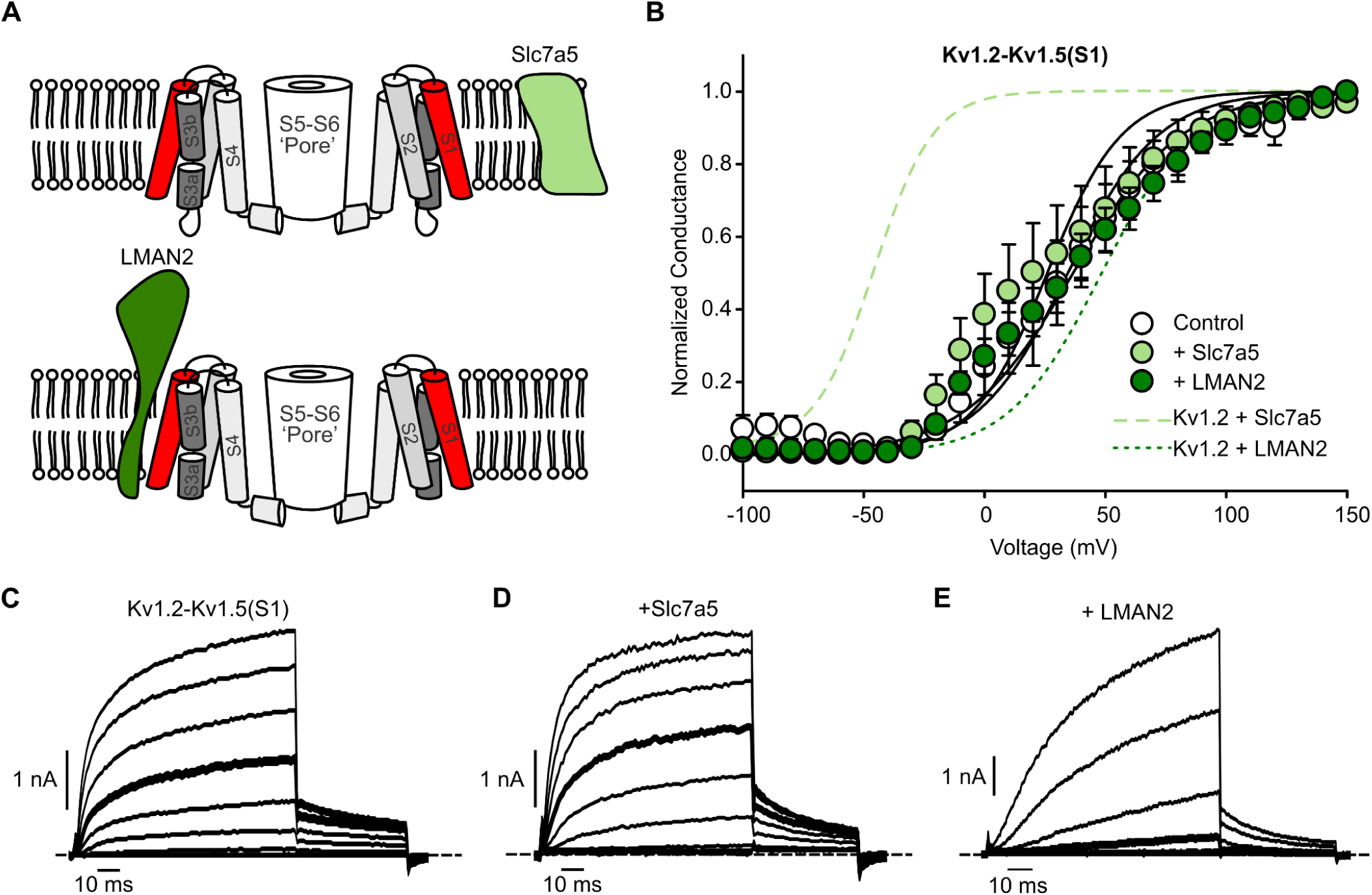
The S1 voltage sensor segment predominantly affects Kv1.2 sensitivity to Slc7a5. (A) Schematic model of Kv1.2-Kv1.5(S1) chimera channel along with candidate interactors Slc7a5 or LMAN2, with chimeric insertion of the Kv1.5 S1 segment highlighted in red. (B) Conductance-voltage relationships for Kv1.2-Kv1.5(S1) co-expressed with Slc7a5 and with LMAN2 as indicated, using an identical protocol as Figure 1. Relationships for Kv1.2 + Slc7a5 (dashed light green) and Kv1.2 + LMAN2 (dashed dark green) are duplicated from Figure 1 for reference. Conductance-voltage fits were (mean ± SEM, Kv1.2-Kv1.5(S1) V_1/2_ = 34.9 ± 9.8 mV, k = 22.0 ± 2.5 mV, n=8; Kv1.2-Kv1.5(S1) + Slc7a5, V_1/2_ = 11.8 ± 6.9, k = 22.7 ± 2.8 mV, n=8; Kv1.2-Kv1.5(S1) + LMAN2 V_1/2_ = 26.2 ± 11.5 mV, k = 21.9 ± 2.0 mV, n=7). (C, D, E) Exemplar patch clamp records from Kv1.2-Kv1.5(S1), Kv1.2-Kv1.5(S1) + Slc7a5 and Kv1.2-Kv1.5(S1) + LMAN2 as indicated. Voltage steps are shown in 20 mV intervals, and pulses to +40 mV are highlighted in boldface for comparison between conditions.

We have previously reported that redox sensitivity of Kv1.2 is strongly influenced by slight amino acid differences in the S2-S3 linker, but the effects of this segment on LMAN2 sensitivity, or the mechanistic relationship between LMAN2 and redox sensitivity, were not firmly established (Baronas *et al*., 2017; Lamothe *et al*., 2024). We observed that replacing the S2-S3 linker of Kv1.2 with Kv1.5 sequence (resulting in two amino acid differences, F251S and T252R), completely abolishes Kv1.2 sensitivity to LMAN2 (Figure 4A-C, E), but preserves sensitivity to Slc7a5. The Kv1.2-Kv1.5(S2-S3L) co-expressed with LMAN2 exhibited a V_1/2_ of -8.8 ± 5.0 mV, indistinguishable from Kv1.2-Kv1.5(S2-S3L) expressed alone (V_1/2_ = -8.4 ± 4.0 mV). In contrast, Kv1.2-Kv1.5(S2-S3L) retained prominent sensitivity to Slc7a5, leading to a hyperpolarizing shift with Slc7a5 (V_1/2_ of -40.9 ± 3.5 mV, Figure 4B,D). The selectivity for Slc7a5 vs. LMAN2 of the Kv1.2-Kv1.5(S2-S3L) chimera is pronounced, relative to the S1 chimera described in Figure 3. Overall, this indicates that the candidate regulators LMAN2 and Slc7a5 rely on distinct channel segments for modulation of function, although they are directly adjacent to one another in the channel sequence, and there may be some overlap.

**Figure 4.**
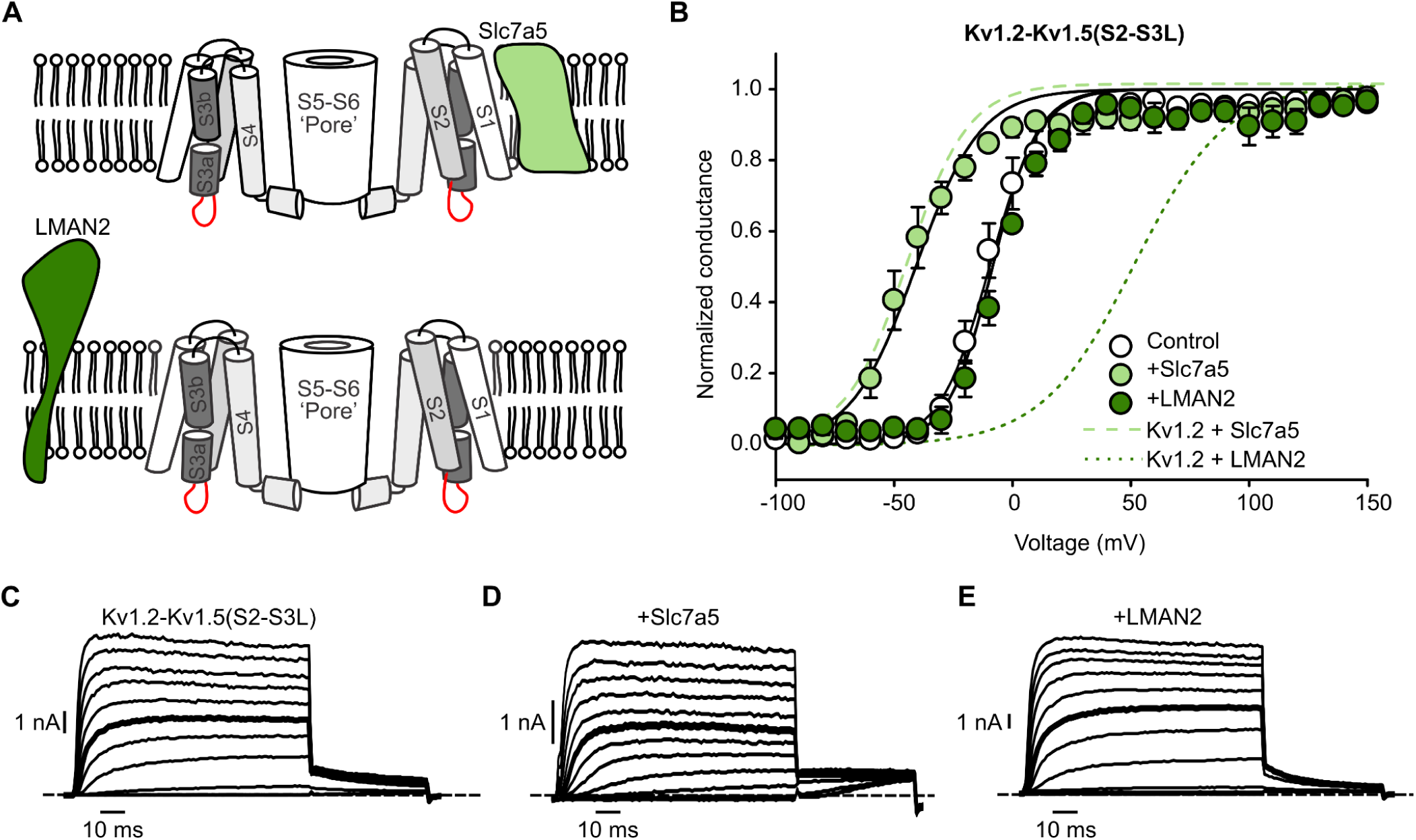
The S2-S3 linker segment predominantly affects Kv1.2 sensitivity to LMAN2. (A) Schematic model of Kv1.2-Kv1.5(S2-S3L) chimera channel along with candidate interactors Slc7a5 or LMAN2, with chimeric insertion of the Kv1.5 S2-S3 linker segment highlighted in red. (B) Conductance-voltage relationships for Kv1.2-Kv1.5(S2-S3L) co-expressed with Slc7a5 or LMAN2 as indicated, using an identical protocol as Figure 1 (dashed lines are duplicated from Figure 1 for reference). Conductance-voltage fits were (mean ± SEM, Kv1.2-Kv1.5(S2-S3L) V_1/2_ = -8.4 ± 4.0 mV, k = 11.0 ± 1.9 mV, n=7; Kv1.2-Kv1.5(S2-S3L) + Slc7a5, V_1/2_ = -40.9 ± 3.5 mV, k = 14.6 ± 2.1 mV, n=5; Kv1.2-Kv1.5(S2-S3L) + LMAN2 V_1/2_ = -8.8 ± 5.0 mV, k = 13.4 ± 2.0 mV, n=5). (C, D, E) Exemplar patch clamp records from Kv1.2-Kv1.5(S2-S3L), Kv1.2-Kv1.5(S2-S3L) + Slc7a5 and Kv1.2-Kv1.5(S2-S3L) + LMAN2 as indicated. Voltage steps are shown in 20 mV intervals, and pulses to +40 mV are highlighted in boldface for comparison between conditions.

### Mutual antagonism of Kv1.2 modulation by Slc7a5 and LMAN2

We tested how the dramatically different functional outcomes of channel modulation by Slc7a5 and LMAN2 might interact or be integrated into channel responses. By co-expressing these modulators, we observed that functional interactions of these regulatory mechanisms can lead to complex patterns of voltage-dependent activation. Firstly, co-expression of Kv1.2 with Slc7a5 and LMAN2 together (Figure 5A) leads to a clear bi-modal voltage-dependence of activation (Figure 5B), which we quantified and fit with a sum of two Boltzmann functions. At negative voltages, there is a fraction of channels that activate with a V_1/2_ of -32.5 ± 3.6 mV, roughly comparable to the V_1/2_ for Slc7a5-modulated Kv1.2.

**Figure 5.**
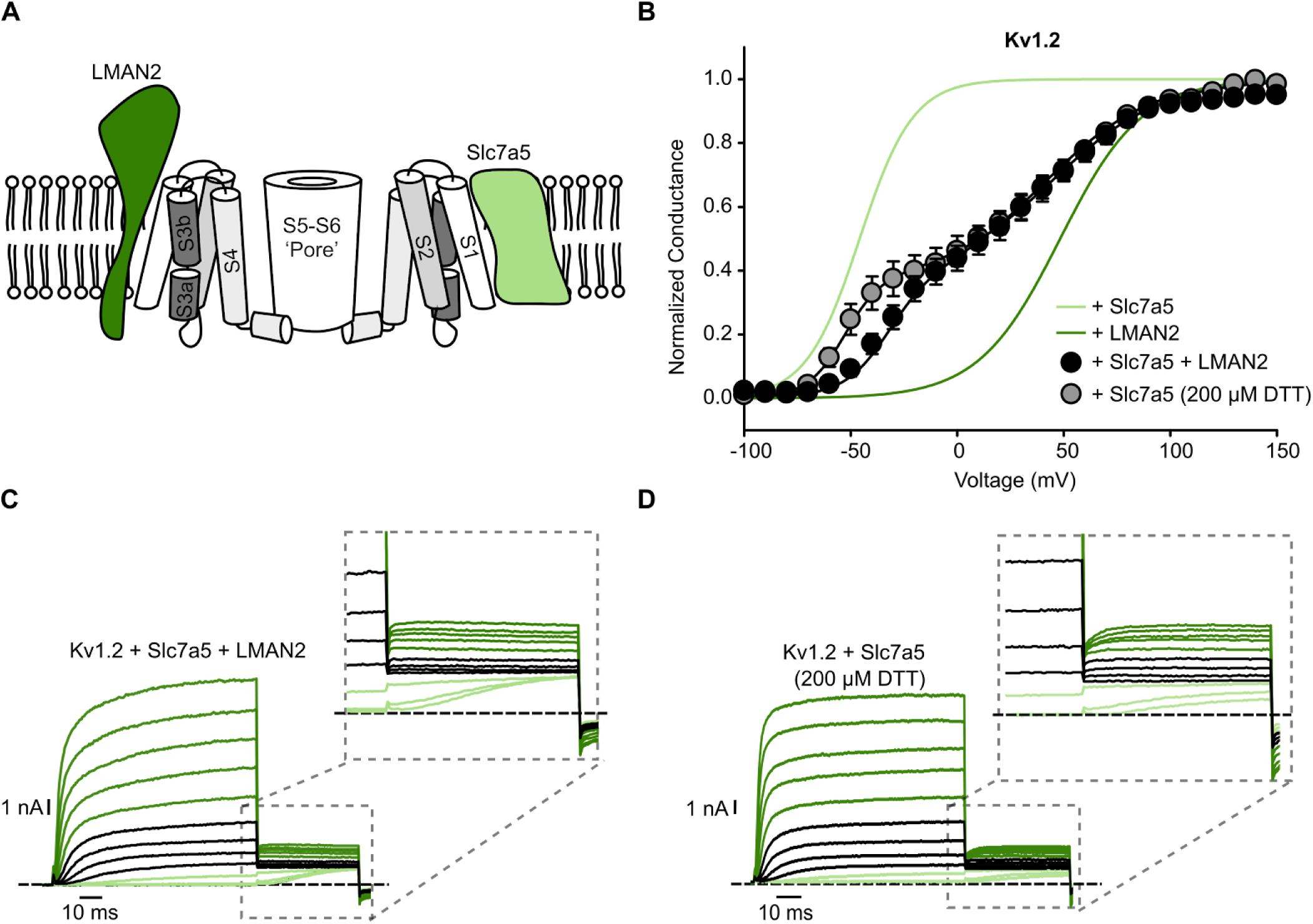
Mutual antagonism of Kv1.2 modulation by Slc7a5 and LMAN2. (A) Schematic model illustrating the potential for modulation by both Slc7a5 and LMAN2. (B) Conductance-voltage relationship for Kv1.2 co-expressed with combinations of Slc7a5 and LMAN2 (black) as well as Kv1.2 co-expressed with Slc7a5 and DTT(200 µM) (gray) as indicated, using an identical protocol as in Figure 1 (dashed lines are duplicated from Figure 1 for reference). Conductance-voltage fits for Kv1.2+Slc7a5+LMAN2 and for Kv1.2+Slc7a5+DTT(200 µM), respectively were (mean ± SEM, Kv1.2+Slc7a5+LMAN2 V_1/2_,_1_ (Slc7a5 component) = -32.5 ± 3.6 mV, k_1_ = 10.0 ± 1.0 mV, V_1/2,2_ (LMAN2 component) = 40.6 ± 6.6 mV, k_2_ = 25.1 ± 2.5 mV, n=14; Kv1.2+Slc7a5+DTT V_1/2_,_1_ (Slc7a5 component) = -52.9 ± 1.2 mV, k_1_ = 8.6 ± 1.3 mV, V_1/2,2_ (DTT component) =44.9 ± 2.7 mV, k_2_ = 25.6 ± 2.3 mV n=6). (C-D) Exemplar patch clamp record from WT Kv1.2 co-expressed with both Slc7a5 and LMAN2 as well as Kv1.2+Slc7a5+DTT respectively, with the inset highlighting the tail currents. Voltage steps are shown in 20 mV intervals for clarity. Pulses from -20 to +40 mV are colored black to highlight voltages that generate the plateau phase of the conductance-voltage relationship. Currents at higher voltages modulated by LMAN2 or DTT are shown in dark green. Currents at lower voltages (modulated by Slc7a5) are shown in light green.

There is a ‘plateau’ region at intermediate voltages, and then a right-shifted component of the curve that activates with a V_1/2_ of +40.6 ± 6.6 mV, comparable to LMAN2-modulated channels. These bi-modal features are clearly apparent in the representative recordings (along with tail current insets, Figure 5C). There is a fraction of total current that exhibits fast activation at hyperpolarized voltages (light green sweeps), a plateau phase that extends across ∼60 mV (black sweeps highlighting -20 mV - +40 mV, inclusive), followed by further activation at more positive voltages (dark green shading, Figure 5C, +60 mV - +140 mV inclusive).

The complex voltage-dependence of gating can also be manipulated with experimental conditions previously shown to promote the slow gating mode (i.e. mimic LMAN2 modulation) (Baronas *et al*., 2017). More specifically, extracellular reducing conditions strongly promote the LMAN2-related slow gating mode of Kv1.2 (with identical requirement for the S2-S3 linker for sensitivity). In terms of complex modulation of Kv1.2 V_1/2_ described thus far, treatment with DTT also mimics the effects of LMAN2 (Figure 5B-D). While Slc7a5 co-expression generates a prominent V_1/2_ shift to negative voltages, application of DTT (200 µM) attenuates this effect and generates a bi-modal voltage-dependence of Kv1.2 gating (Figure 5B,D). We found that this attenuation of the Slc7a5-mediated gating component could be exaggerated even further by co-expression/treatment with increasing amounts of LMAN2 and DTT together (Figure 6). This demonstrates a progression of attenuation of the Slc7a5-mediated gating shift, as we increased the extent of the counteracting LMAN2-related mechanism. Overall, these findings illustrate that the influence of multiple modulators related to Slc7a5 and LMAN2/DTT can functionally interact and generate complex patterns of voltage-dependent activation of Kv1.2, spanning a wide dynamic range of voltage. It seems reasonable to think of this as a ‘competitive’ type of interaction, because the conductance-voltage relationships are quite clearly resolved into two distinct components. This indicates that there are populations of channels influenced by LMAN2, and Slc7a5, but not both.

**Figure 6.**
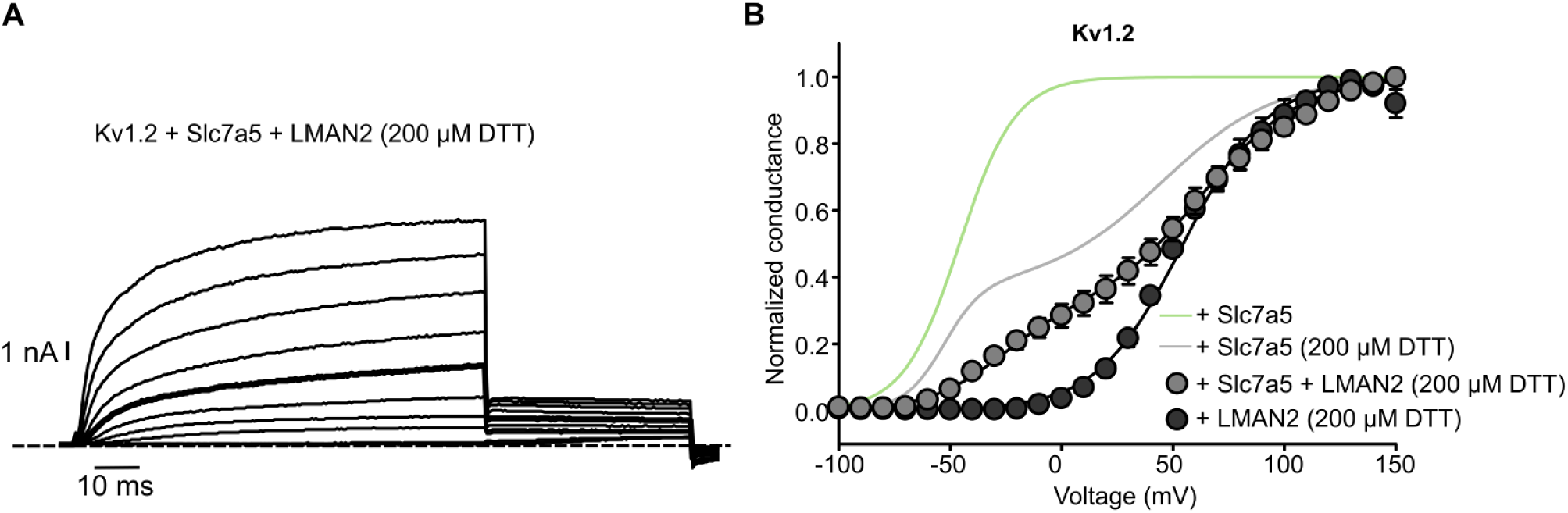
LMAN2 and DTT further attenuate Slc7a5-mediated gating effects on Kv1.2. (A) Exemplar patch clamp records from Kv1.2 + Slc7a5 + LMAN2 + DTT. Voltage steps are shown in 20 mV intervals for clarity. Pulses to +40 mV have been highlighted in boldface for comparison between conditions. (B) Conductance voltage-relationship for Kv1.2 co-expressed with Slc7a5 and LMAN2 +DTT(200 µM) as well as LMAN2 and DTT(200 µM) as indicated, using an identical protocol as Figure 1 (For reference, green and gray lines are duplicated from Figure 1 and Figure 5, respectively for reference). Conductance-voltage fit was (mean ± SEM, Kv1.2+Slc7a5+LMAN2+DTT V_1/2_,_1_ (Slc7a5 component) = -27.6 ± 6.2 mV, k_1_ = 16.6 ± 2.8 mV, V_1/2,2_ (LMAN2 and DTT component) = 63.5 ± 4.4 mV, k_2_ = 22.1 ± 1.1 mV n=16; Kv1.2+LMAN2+DTT V_1/2_ =54.6 ± 2.5 mV, k_1_ = 19.7 ± 2.0 mV, n=4).

An important consideration is whether the antagonism of Slc7a5 effects by LMAN2 and/or DTT is related to competition for actions on the channel, or a direct effect of LMAN2/DTT on Slc7a5. We tested these possibilities by investigating the effects of Slc7a5 and LMAN2 (or DTT) co-expression on the Kv1.2-Kv1.5(S2-S3L) chimera, which is wholly insensitive to DTT and LMAN2 (shown in Figure 4). We observed that LMAN2/DTT has only minimal effects on the Slc7a5-mediated gating shift, indicating that LMAN2/DTT actions on the channel are essential for influencing Slc7a5 effects (Figure 7A). Therefore, it is most likely that the bi-modal gating properties (shown in Figure 5) and attenuation of Slc7a5-mediated gating shifts (Figure 6) arise from competitive actions of LMAN2 and Slc7a5 on the channel, because the S2-S3L chimera exhibits no attenuation when treated with DTT.

**Figure 7.**
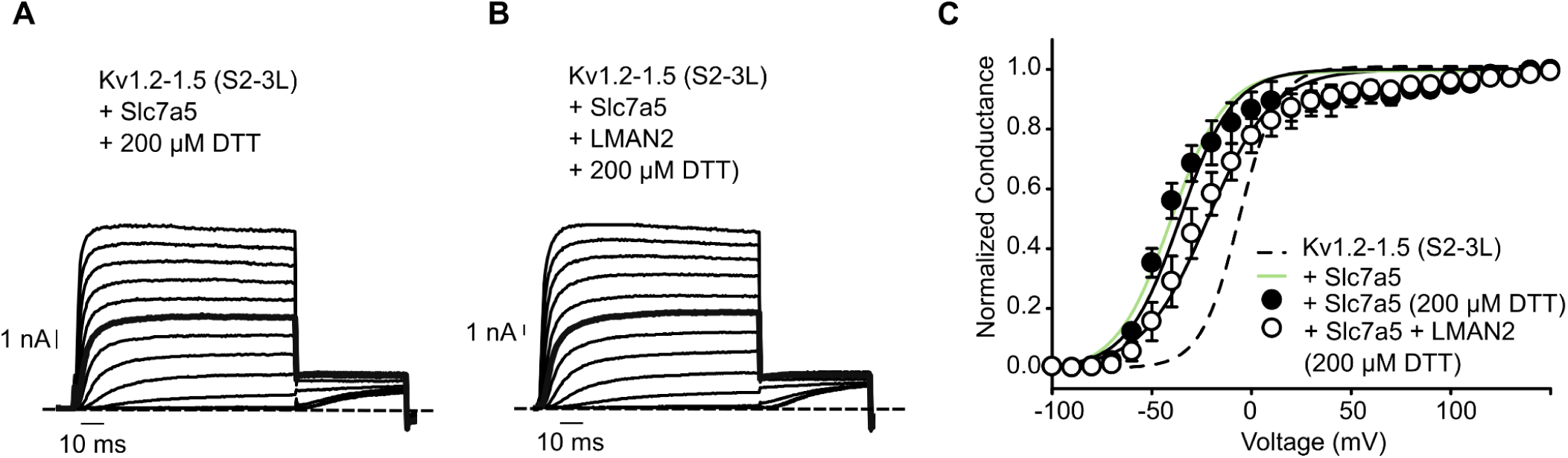
Antagonistic effects of LMAN2/DTT on Slc7a5 are due to direct actions on Kv1.2. Representative traces and conductance-voltage relationships for Kv1.2-Kv1.5(S2-S3L) co-expressed with Slc7a5 and treated with DTT (200 µM) (A),(C, black circles) or Slc7a5 + LMAN2 and DTT (200 µM) (B),(C, white circles) as indicated, using an identical protocol as Figure 1. Relationships for Kv1.2-Kv1.5(S2-S3L) (dashed black fit) and Kv1.2-Kv1.5(S2-S3L) + Slc7a5 (light green) are duplicated from Figure 3 for reference. Conductance-voltage fits were (mean ± SEM, Kv1.2-Kv1.5(S2-S3L) + Slc7a5 + DTT V_1/2_ = -35.9 ± 6.1 mV, k = 16.6 ± 5.5 mV, n=4; Kv1.2-Kv1.5(S2-S3L) + Slc7a5 + LMAN2 + DTT V_1/2_ = -21.9 ± 6.2 mV, k = 17.7 ± 2.6 mV, n=5).

Related to this finding, bi-modal features are altered in a predictable modulator specific manner in each of the various Kv1.2-Kv1.5 chimeric channels described thus far (Figure S1). The S2-S3L chimera (which selectively ablates LMAN2-sensitive modulation), exhibits selective loss of the LMAN2-mediated component when cells are co-transfected with both LMAN2 and Slc7a5, and thus channels gate uniformly with a strong hyperpolarized V_1/2_ (V_1/2_ = -32.8 ± 4.5 mV) (Figure S1B,E). In contrast, the Kv1.2-Kv1.5(S1) chimera (which markedly weakens the effects of Slc7a5), exhibits the LMAN2-mediated component when transfected with both Slc7a5 and LMAN2, exemplified by slow activation kinetics and a V_1/2_ that is prominently shifted to positive depolarized voltages (V_1/2_ = 26.6 ± 9.4 mV) (Figure S1C,F). As mentioned previously, there are some uncertainties with interpretation of data from the Kv1.2-Kv1.5(S1) chimera, as there is a gating shift to positive voltages, even in the absence of co-transfected LMAN2 (Figure 3, Figure S1C). This is likely due to some basal levels of modulation in these cells in the absence of exogenous LMAN2 (seen in both Figure 3 and S1C). It is also possible that the S1 segment influences LMAN2 modulation (although the S2-S3 linker is clearly the most important determinant), as the maximal LMAN2-mediated gating shift in the S1 chimera is not as large as in WT Kv1.2 channels (Figure 3B). Lastly, the Kv1.2-Kv1.5(S1-S3) chimera exhibited essentially no modulation when co-expressed with Slc7a5 and LMAN2 together (V_1/2_ = 5.5 ± 4.7 mV) (Figure S1A,D).

### Correlation of gating properties with relative expression of Slc7a5 and LMAN2

We emphasize that the bi-modal G-V curve described in Figure 5 (arising with co-expression of Slc7a5 and LMAN2) reflects the average of recordings from multiple cells. However, the gating properties observed in individual cells can deviate significantly depending on relative amounts of LMAN2 or Slc7a5 (leading to relatively different contributions of each component on the conductance-voltage relationship). That is, in some cells there is a large component with a negative V_1/2_ (attributed to Slc7a5), whereas in other cells this component may be smaller. To validate this finding, we used a multi band-pass filter to assess the relative expression levels of both LMAN2 and Slc7a5 prior to patch clamping each cell (Figure 8A). Cells were co-transfected with mCherry-Slc7a5 and EGFP-LMAN2, and a sample image viewed through the multi bandpass filter is shown in Figure 8A. Varying patterns of expression can be seen in cells that are strongly red (predominantly expressing Slc7a5), strongly green (predominantly expressing EGFP-LMAN2), or yellow (some level of expression of both modulators). Individual G-V curves from cells recorded in these conditions are shown in Figure 8B, and color coded based on the predominant color seen through the multi bandpass filter. It is apparent that bi-modal gating is seen frequently in yellow cells (with both Slc7a5 and EGFP), whereas red vs. green cells are biased towards Slc7a5-like or LMAN2-like modulation, respectively. These findings further support the notion that multimodal gating of Kv1.2 arises from distinct Slc7a5- and LMAN2-mediated components.

**Figure 8.**
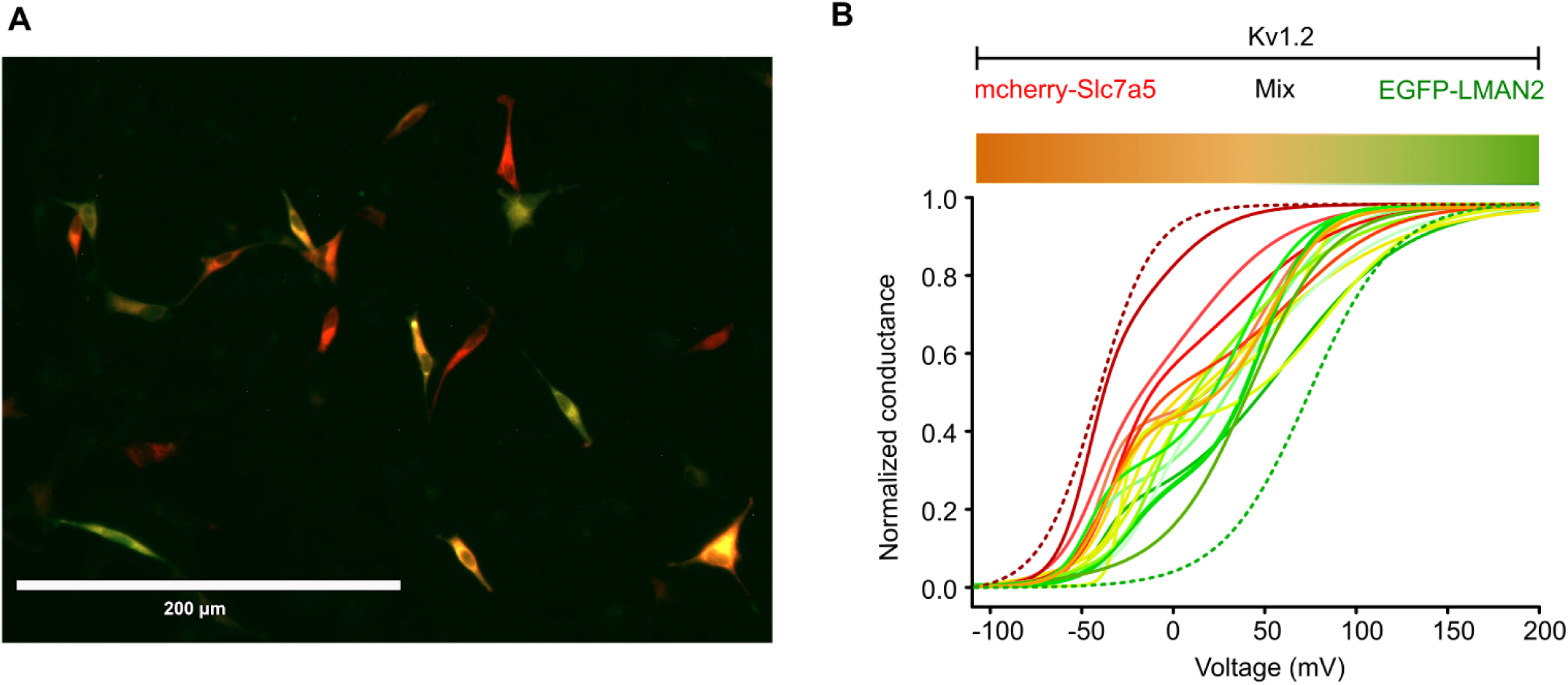
Relative expression of Slc7a5 and LMAN2 predicts gating modulation of Kv1.2. (A) 20X microscopic field of view of visible color ranges from cells expressing differing amounts of mcherry-Slc7a5 and/or EGFP-LMAN2. (B) Color gradient denoting how the relative expression levels of mcherry-Slc7a5 (red) and EGFP-LMAN2 (green) (as seen under the microscope) affect the conductance-voltage relationship in WTKv1.2 channels from individual cells. Fits for Kv1.2 + Slc7a5 (dotted red) and Kv1.2 + LMAN2 (dotted green) are duplicated from Figure 1 for reference.

### Summarized overview of gating modulation in Kv1.2-Kv1.5 chimeric channels

The gating effects of Slc7a5 and LMAN2 on all chimeras tested is summarized in Figure 9. Key features of this summarized data are that co-expression of either Slc7a5 or LMAN2 has dramatically different effects on WT Kv1.2 gating. Co-expression of WT Kv1.2 with both Slc7a5 and LMAN2 simultaneously generates a bi-modal voltage-dependence that can be fit with two separate components that are roughly similar to the gating shifts generated by Slc7a5 or LMAN2 alone. Various chimeras can be engineered to weaken all modulation (in the S1-S3 chimera), selective weakening of LMAN2 modulation (in the S2-S3L chimera), or quasi-selective weakening of Slc7a5 modulation (in the S1 chimera). Overall, these findings indicate that complex patterns of voltage-dependence emerge when Kv1.2 channels are co-expressed with LMAN2 and Slc7a5, likely due to mutual antagonism of the gating modulation caused by Slc7a5 or LMAN2.

**Figure 9.**
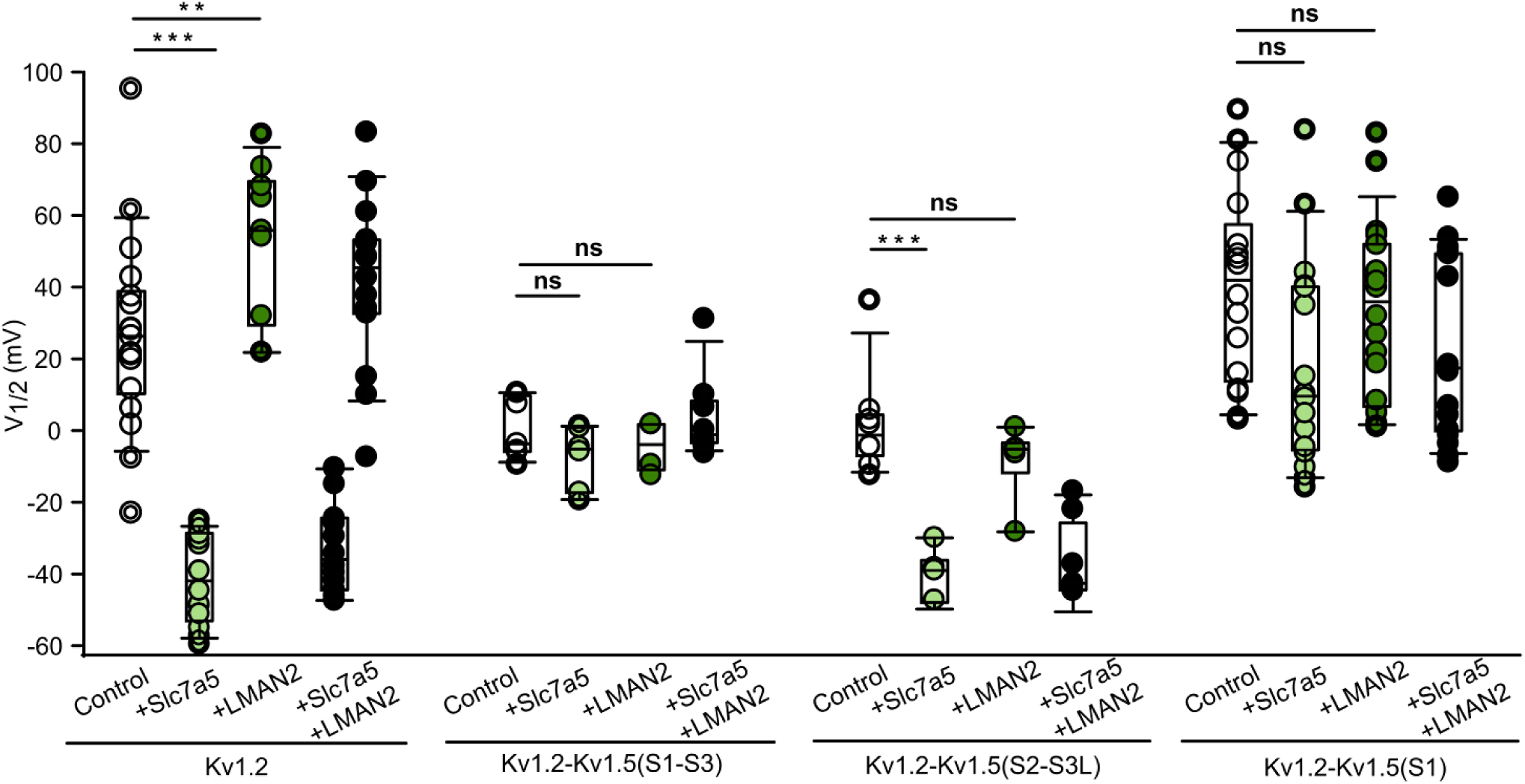
Summary of the effects of LMAN2 and Slc7a5 on the V_1/2_ of Kv1.2-Kv1.5 chimeric channels. Individual data points of the voltage of half-activation of WT Kv1.2 and Kv1.2-Kv1.5 chimeras in control (white), Slc7a5 alone (light green), LMAN2 alone (dark green) and co-expression of both modulators (black). A One-way ANOVA followed by Dunnett’s post-hoc test was used to compare Slc7a5 or LMAN2 conditions vs. Control within each channel type. P-values are denoted above the box plots, where *** denotes a p-value <0.001, ** denotes a p-value <0.01 and ‘ns’ is statistically non-significant.

### Distinct effects of Slc7a5 on gating and expression of Kv1.2

We previously reported that Slc7a5 weakens overall expression and maturation when co-expressed with Kv1.2 (Baronas *et al*., 2018; Lamothe and Kurata, 2020). This is apparent as a change in protein abundance on Western blot, along with weaker appearance of bands corresponding to mature glycosylated forms of Kv1.2 in the molecular weight range around 75 kDa (Figure 10). It has been unclear whether this effect on expression is related to the gating effects discussed thus far, or whether there may be different underlying signaling pathways or mechanisms. We tested the expression of each of the chimeras used in this study (S1, S2-S3L, S1-S3), when co-expressed with Slc7a5, LMAN2, or both. We observed that Slc7a5 attenuated expression and maturation in all of the constructs tested, even the S1 chimera that is resistant to Slc7a5-mediated gating effects. LMAN2 did not markedly affect expression levels, while co-expression of Slc7a5 and LMAN2 together resembled the effects of Slc7a5. Thus there appears to be a difference in the molecular determinants that govern the sensitivity to Slc7a5 gating modulation versus expression and maturation.

**Figure 10.**
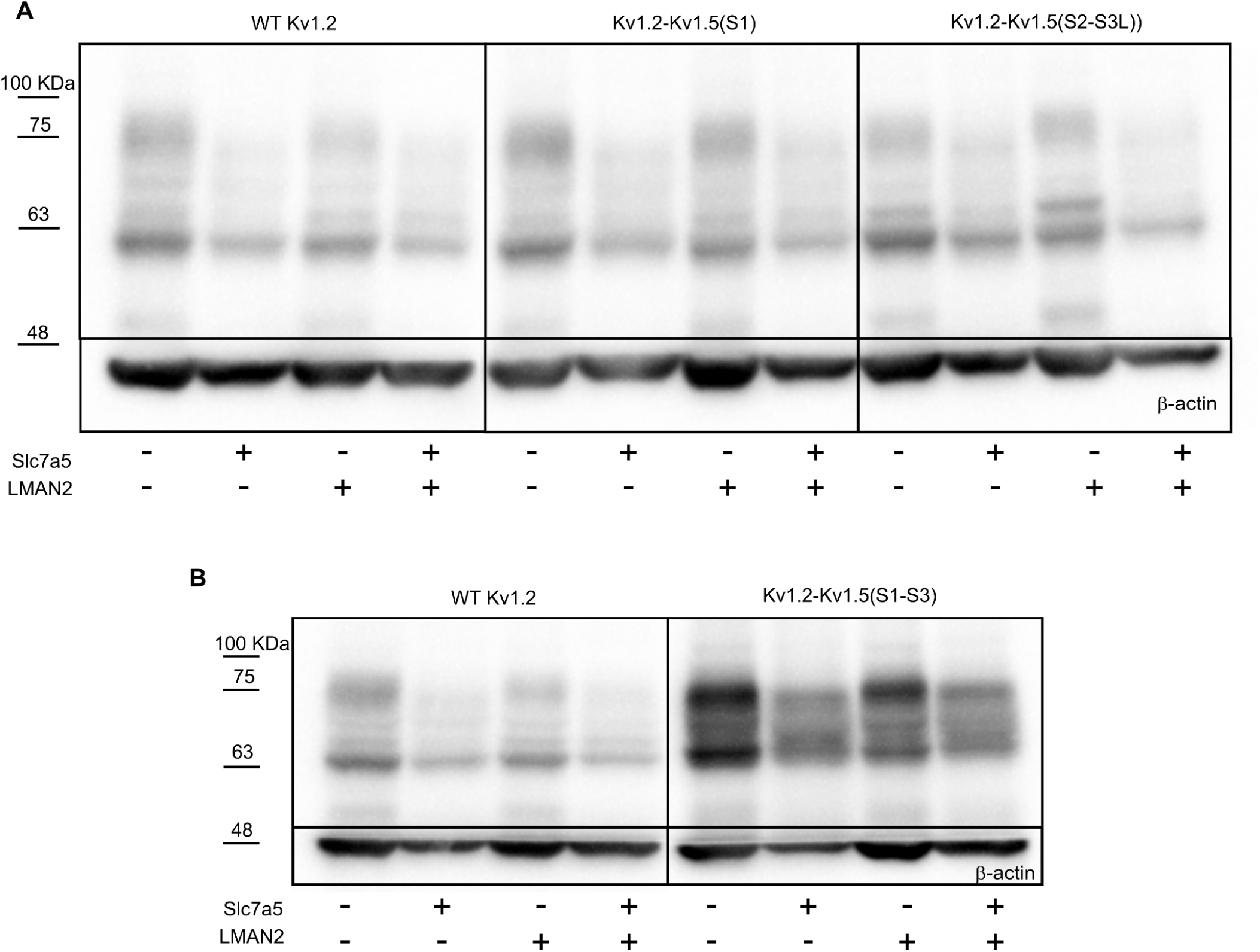
Slc7a5 alters the expression and maturation of Kv1.2:Kv1.5 chimeras. (A) Exemplar anti-Kv1.2 immunoblot of protein lysate from HEK293 cells transfected with WT Kv1.2, Kv1.2+Slc7a5 (1:1), Kv1.2+LMAN2 (1:1) and Kv1.2+Slc7a5+LMAN2 (1:1:1) as well as corresponding transfections for Kv1.2-Kv1.5 (S1) and (S2-S3L) chimeras, respectively, for 72 h. □-actin was used as a loading control. Exemplar anti-Kv1.2 immunoblot of protein lysate from HEK293 cells transfected with WT Kv1.2 or Kv1.2-Kv1.5(S1-S3), +Slc7a5, +LMAN2 and +Slc7a5+LMAN2 under similar conditions as (A).

## DISCUSSION

Kv1.2 exhibits unexplained variability of its gating properties when characterized in both heterologous and primary cell systems (Grissmer *et al*., 1994; Rezazadeh *et al*., 2007; Baronas, McGuinness, *et al*., 2015; Baronas *et al*., 2017; Abraham *et al*., 2019; Lamothe *et al*., 2020). Even in heterologous systems where channels can be faithfully recorded with minimal contributions from other endogenous channels, Kv1.2 exhibits exceptionally wide variation of the voltage-dependence of activation between cells, and inconsistent fitting by Boltzmann functions that are usually adequate to describe most Kv channel behavior (Baronas, Yang, *et al*., 2015; Lamothe *et al*., 2020). This seems to indicate prominent regulation by cellular factors that are not explicitly controlled in these experiments, and past work has highlighted numerous proteins or factors like extracellular redox potential, that regulate gating and/or expression of Kv1.2 and related Kv1 channels (Kourrich *et al*., 2013; Baronas *et al*., 2018; Abraham *et al*., 2019; Lamothe *et al*., 2024). However, there is little understanding of how these various regulators may functionally interact, and also what structural elements of Kv1.2 enable the influence of diverse regulatory proteins.

We have worked towards understanding proteins and signaling mechanisms that may influence Kv1.2 gating. In this study, we focused on two membrane proteins (Slc7a5 and LMAN2) that are particularly interesting because of the prominent shifts in voltage-dependent gating that they elicit, highlighting an extraordinary dynamic range of voltages across which Kv1.2 can be regulated. Slc7a5 and LMAN2 promote two extremes of voltage-dependent gating - Slc7a5 shifts the V_1/2_ of Kv1.2 prominently to negative voltages, in the range of -50 mV, whereas LMAN2 markedly decelerates activation and causes a prominent shift to positive voltages (Baronas *et al*., 2018; Lamothe *et al*., 2020, 2024). Under ambient conditions, when Kv1.2 is transfected without any additional exogenous candidate regulators, its cell to cell behavior is ‘bounded’ within these two extremes, and so we considered the contributions of LMAN2 and Slc7a5 in terms of potential mechanisms underlying cell to cell variability of Kv1.2 gating.

Our findings indicate that the voltage-sensing domain is a prominent structural element that influences Kv1.2 sensitivity to regulatory proteins. Sensitivity to both LMAN2 and Slc7a5 is completely attenuated with substitution of the S1-S3 segments of Kv1.5. This is consistent overall with our prior investigations of Slc7a5 modulation and redox sensitivity, which used a similar strategy of Kv1.2/Kv1.5 chimeric channels to identify critical structural determinants of regulation (Lamothe *et al*., 2020, 2024). It is noteworthy that the Kv1 channels exhibit a high degree of homology/identity within the transmembrane segments, with most sequence differences concentrated within the N- and C-termini, along with the extracellular S1-S2 linker. Thus, we have found it interesting and surprising that structural elements underlying the subtype specific effects of Slc7a5 and LMAN2 are located in regions of high sequence conservation. Slc7a5 modulation of activation is most prominently controlled by the S1 segment, and can be significantly attenuated by the Kv1.2[I164A] mutation (Lamothe *et al*., 2020). The intracellular S2-S3 linker is the most important determinant of sensitivity to LMAN2 and redox, and these are completely abolished by the Kv1.2[F251S][T252R] double mutation (Lamothe *et al*., 2024).

Investigation of mechanisms of modulation of Kv1 channels has primarily focused on cytoplasmic signaling proteins and post-translational modifications of the N- and C-termini. The most well characterized regulatory partners are the Kvβ subunits, which interact with the ordered N-terminal T1 domain (Rettig *et al*., 1994; Heinemann *et al*., 1996; Gulbis *et al*., 2000). Other proposed regulatory mechanisms include kinase modulation at phosphosites in the N- and C-termini, and association with various scaffolding/cytoskeletal proteins that may influence maturation or localization of channels (Hattan *et al*., 2002; Yang *et al*., 2007; Connors *et al*., 2008; Ogawa *et al*., 2008; Sanders *et al*., 2020). There is growing consideration of Kv channel association and regulation by transmembrane proteins like LMAN2 and Slc7a5. For example, Kv4 channels assemble with the single pass transmembrane protein DPPX, which influences localization and inactivation kinetics (Jerng *et al*., 2005; Amarillo *et al*., 2008). BK channels have a slightly different architecture of the voltage-sensing domain (comprising an additional transmembrane segment), but also associate with transmembrane β- and γ-subunits with powerful modulatory effects (Gonzalez-Perez *et al*., 2015; Gonzalez-Perez and Lingle, 2019). In the case of Kv1.2 or other Kv1 channels, there is recognition of multiple transmembrane proteins with potential effects on gating or localization, including OPRS1, Slc7a5, LMAN2, and the LGI-ADAM complex, although ongoing investigation will help develop a consensus on the cell types and physiological systems where these potential modulators are most important (Baronas *et al*., 2018; Abraham *et al*., 2019; Lamothe and Kurata, 2020; Fukata *et al*., 2021; Kozar-Gillan *et al*., 2023; Lamothe *et al*., 2024; Zhou *et al*., 2024). Our recent findings have demonstrated that native Kv1.2 currents in hippocampal neurons and DRG neurons exhibit regulation that resembles the effects of LMAN2 and redox reported here and elsewhere (Baronas, McGuinness, *et al*., 2015; Lamothe *et al*., 2024).

The limited space and surface for interactions in the VSD will likely influence how various proteins affect channel function when expressed together. In this study for example, our findings illustrate that Slc7a5 effects are mutually exclusive from the gating effects of LMAN2 or extracellular reducing conditions.

This is most clearly shown in Figure 5, where multiple components of voltage-dependent activation are clearly resolvable in the conductance-voltage relationships. Our interpretation of this observation is that the ensemble of channels in the cell comprises distinct populations that are regulated by either Slc7a5 or LMAN2/redox. From a physical perspective, we presume that there are constraints that prevent both proteins from associating with Kv1.2 at once, and this may explain the distinct activation components. It is noteworthy that the vast difference in voltage-dependence of activation between these conditions allows us to resolve distinct components, but we can imagine that other regulator ratios could blunt the conductance-voltage relationships (leading to an apparent shallow G-V). This is also different from the regulation of BK channels by transmembrane β- and γ-subunits, which can both co-assemble and simultaneously influence channel function (Gonzalez-Perez *et al*., 2015). Overall, the competitive interactions of Slc7a5 or LMAN2 with Kv1.2 seems to underlie the generation of complex multi-component conductance-voltage relationships.

In summary, our findings demonstrate that multiple regulatory proteins converge on the Kv1.2 voltage-sensing domain, with powerful effects on gating and expression. Effects on channel function appear to be competitive, in the sense that co-expression with Slc7a5 and LMAN2 generates complex conductance-voltage relationships with components clearly attributable to Slc7a5 or LMAN2. These findings highlight the extraordinarily wide range of voltage-dependent gating modulation that can be achieved for Kv1.2 channels. We do not observe remotely similar levels of modulation of other closely related Kv1 channels, and so it appears that this feature is highly subtype specific and governed by elements of the voltage-sensing domain.

## SUPPLEMENTARY FIGURES

**Supplementary Figure S1.**
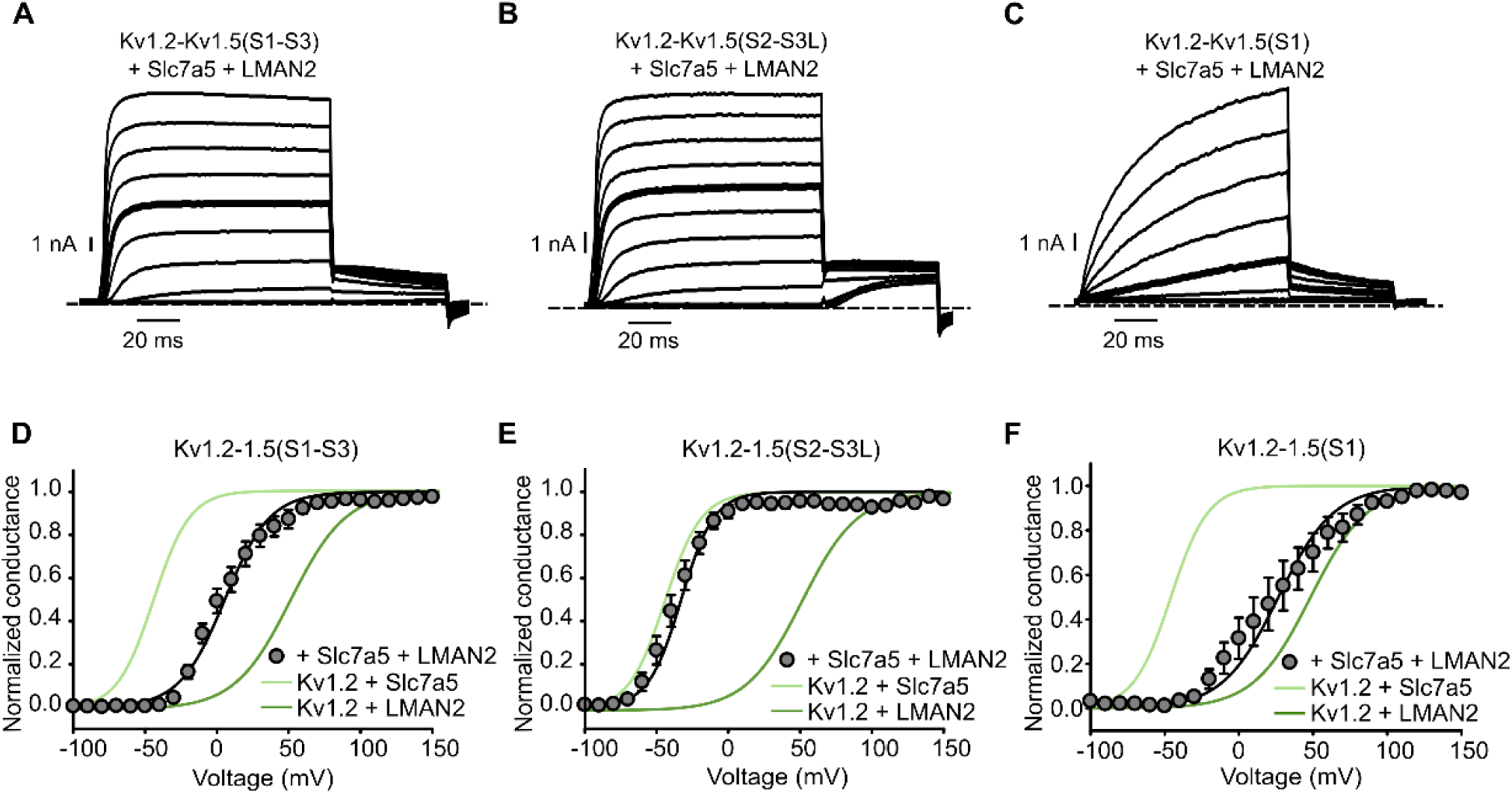
Selective attenuation of LMAN2 or Slc7a5 modulation of Kv1.2:Kv1.5 chimeric channels. (A,B,C) Exemplar patch clamp records from each Kv1.2-Kv1.5 chimeric channel co-expressed with both Slc7a5 and LMAN2. Voltage steps are shown in 20 mV intervals, and pulses to +40 mV are highlighted in boldface for comparison between conditions. (D,E,F) Conductance voltage-relationships for indicated Kv1.2-Kv1.5 chimeric channels co-expressed with both Slc7a5 and LMAN2. Relationships for Kv1.2+Slc7a5 (light green line), and Kv1.2+LMAN2 (dark green line) are duplicated from Figure 1 for reference. Conductance-voltage fits were (mean ± SEM, Kv1.2-Kv1.5(S1-S3) + Slc7a5 + LMAN2 V_1/2_ = 5.5 ± 4.7 mV, k = 16.3 ± 1.3 mV, n=7; Kv1.2-Kv1.5(S2-S3L) + Slc7a5 + LMAN2 V_1/2_ = -32.8 ± 4.5 mV, k = 13.8 ± 1.9 mV, n=9; Kv1.2-Kv1.5(S1) + Slc7a5 + LMAN2 V_1/2_ = 26.6 ± 9.4 mV, k = 20.4 ± 2.3 mV, n=8).

## Notes

### Competing Interest Statement

The authors have declared no competing interest.

